# Integration of visual motion and pursuit signals in areas V3A and V6+ across cortical depth using 9.4T fMRI

**DOI:** 10.1101/2021.12.09.471881

**Authors:** Fatemeh Molaei-Vaneghi, Natalia Zaretskaya, Tim van Mourik, Jonas Bause, Klaus Scheffler, Andreas Bartels

## Abstract

Neural mechanisms underlying a stable perception of the world during pursuit eye movements are not fully understood. Both, perceptual stability as well as perception of real (i.e. objective) motion are the product of integration between motion signals on the retina and efference copies of eye movements. Human areas V3A and V6 have previously been shown to have strong objective (‘real’) motion responses. Here we used high-resolution laminar fMRI at ultra-high magnetic field (9.4T) in human subjects to examine motion integration across cortical depths in these areas. We found an increased preference for objective motion in areas V3A and V6+ i.e. V6 and possibly V6A towards the upper layers. When laminar responses were detrended to remove the upper-layer bias present in all responses, we found a unique, condition-specific laminar profile in V6+, showing reduced mid-layer responses for retinal motion only. The results provide evidence for differential, motion-type dependent laminar processing in area V6+. Mechanistically, the mid-layer dip suggests a special contribution of retinal motion to integration, either in the form of a subtractive (inhibitory) mid-layer input, or in the form of feedback into extragranular or infragranular layers. The results show that differential laminar signals can be measured in high-level motion areas in the human occipitoparietal cortex, opening the prospect of new mechanistic insights using non-invasive brain imaging.

**Significance Statement:** Visual stability and our ability to differentiate between self-induced and real motion are central to our visual sense. Both require the integration of two signals – retinal motion and copies of muscle commands used for eye movements (efference copies). A reasonable assumption is that either the efference copy or the result of integration will be conveyed to high-level visual regions along with visual retinal input, possibly differentially across cortical depth as the input sources differ. Our ultra-high field recordings present the first laminar evidence of differential signal processing of retinal and objective motion signals in area V6+, and present a first window into a mechanistic understanding of visual high-level motion processing.

## INTRODUCTION

Visual motion perception, contrary to our intuition, is only partly based on visual input. Non-visual cues such as efference copies from eye movements (i.e. copies of eye-movement motor commands) that are sent to the visual system determine to an equal degree our perception of motion, or the lack of it. Only the multi-modal integration of visual motion signals with non-visual cues allows for a stable perception of the world and differentiation between self-induced motion and objective (or real) motion (von Holst and Mittelstaedt, 1950; Gibson, 1954; Royden et al., 1992). Patients with selective impairment of such integration abilities point to a parieto-occipital involvement in this process (Haarmeier et al., 1997). Correspondingly, invasive electrophysiology in macaques has revealed several parieto-occipital regions containing varying fractions of so-called ‘real motion’ neurons whose response reflects motion in the environment even when it is cancelled on the retina through pursuit (Galletti et al., 1984, 1988, 1990; Galletti and Battaglini, 1989; Erickson and Thier, 1991; Ilg et al., 2004). Such neurons were predominantly found in motion processing regions V3A, V6, MST, VIP and VPS, but were also present as early as V1 (Galletti et al., 1984, 1988; Zhang et al., 2004; Dicke et al., 2008; Daddaoua et al., 2014).

In the human brain, only few studies addressed the integration of pursuit eye-movements with retinal motion (Arnoldussen et al., 2011; Fischer et al., 2012; Nau et al., 2018). These studies showed that V3A and V6 stand out in a unique way, with overwhelming responses to objective (‘real’) motion, and absent or negative responses to retinal motion (Fischer et al., 2012). Saccade-related spatial updating led to anatomically overlapping activity (Merriam et al., 2003, 2007), and a voxel-wise segregation of these two motion types has been suggested (Arnoldussen et al., 2011). Here, we investigated whether retinal, objective, and pursuit motion signals are segregated across the cortical depth.

Laminar organization of neuronal circuitry in the cortex involves feedforward, lateral, and feedback pathways (Larkum, 2013; Markov et al., 2014), whereby feedback signals enter in deep and superficial layers, and feedforward signals in middle layers (Wong-Riley, 1978; Harris and Mrsic-Flogel, 2013). V3A and V6 have reciprocal connectivity with subcortical as well as cortical regions in both dorsal and ventral streams (Galletti et al., 1990, 2001; Anderson and Martin, 2005), and in particular with parietal cortex and the smooth pursuit region of the frontal eye fields (Stanton et al., 2005). We hypothesized that the latter connections convey eye movement related signals to the feedback layers, and that the retinal motion signals are conveyed to the feedforward middle layers of V3A and V6+. Also, if the previously observed negative contribution of retinal motion in V6+ reflects inhibitory input (Fischer et al., 2012), it would be expected in feedforward middle layers. To test these hypotheses, we used high-resolution fMRI at ultra-high field (9.4T) to quantify neural signals across cortical depth related to retinal motion, objective motion, and pursuit in areas V3A, V6+, and, as a control, in area IPS-0 / V7 (Tootell et al., 1998).

Linking neural computations of a single layer, or column, to the BOLD signal requires imaging with high spatio-temporal resolution. Imaging at high field (> 7T) increases sensitivity of the BOLD signal to the microvasculature (Uludag et al., 2009) resulting in its increased accuracy with respect to the site of neuronal activity. High resolution imaging furthermore reduces contamination of functional signal by physiological noise (Triantafyllou et al., 2005) and decreases contribution of large draining vessels (Olman et al., 2007; Polimeni et al., 2010; Baez-Yanez et al., 2017). Even though recent ultra-high field fMRI studies have provided depth-(Muckli et al., 2015; Kok et al., 2016; Huber et al., 2017; Trampel et al., 2017; Kashyap et al., 2018; Klein et al., 2018) and column-resolved functional signals (Yacoub et al., 2008; Shmuel et al., 2010; Nasr et al., 2016), the physiological limits for laminar imaging are still actively debated (Polimeni et al., 2018).

## METHODS

### Participants

Eleven neurologically healthy adults (3 females, 8 males, mean age 32 years ± 9 SD) with normal or corrected-to-normal vision volunteered to participate in the study. In accordance with the local research ethics committee requirements volunteers underwent a physical and psychological check-up by a local physician and provided written informed consent. All investigations were conducted in agreement with the Declaration of Helsinki and were approved by the ethics committee of the university clinics Tübingen. Prior to scanning, subjects were instructed on the experimental procedures and performed a test trial to get accustomed to stimuli and to the task.

### Visual Stimulation and Experimental Design

Visual stimuli and paradigm (Figure 1) closely replicated those used previously to identify and reliably separate neuronal signals related to retinal and objective (‘real’) motion in V3A and V6 (Fischer et al., 2012). In brief, stimuli consisted of randomly arranged black and white dots (size ranging from 0.1 to 1.1 deg) on a grey (90 cd/m^2^) background, presented at 100% contrast (i.e. maximal luminance for white dots and minimal luminance for black dots). The 320 visible dots yielded an average density of 0.75 dots/deg^2^. The experiment included 4 conditions arranged in a 2×2 factorial design including 2 factors with 2 levels each. The two factors were “pursuit” (on/off) and “objective motion” (on/off). Objective motion was achieved by displacement of the entire dot field along the vertical and horizontal axes along a 2D (i.e. planar) sinusoidal trajectory with either 3 or 4 cycles per trial (randomly assigned to x and y axes, respectively) and with random initial phases. This led to figure-of-eight-style planar trajectories of the dot field. We refer to this as objective motion (equivalent to ‘real’ motion) to distinguish it from retinal motion that can be induced by eye movements over the static dot field. Pursuit was implemented by moving the otherwise centrally presented fixation disc (that contained the fixation task, see Fixation Task below) along the same trajectory. The maximal eccentricity of the fixation disc reached 2.5 visual degrees, such that the eccentricity of the visually fully controlled stimulus (i.e. fixation to screen border) was at least 12.5 × 7.5 visual degrees or more at all times (given the screen size of 30 × 20 visual degrees). When both pursuit and objective motion were ‘on’, the two were coupled, such that the fixation task moved together with the dots, resulting in zero retinal motion. The starting direction was randomized for each trial. The mean speed for objective motion (and pursuit) was 3.80 deg/s.

**Figure 1:**
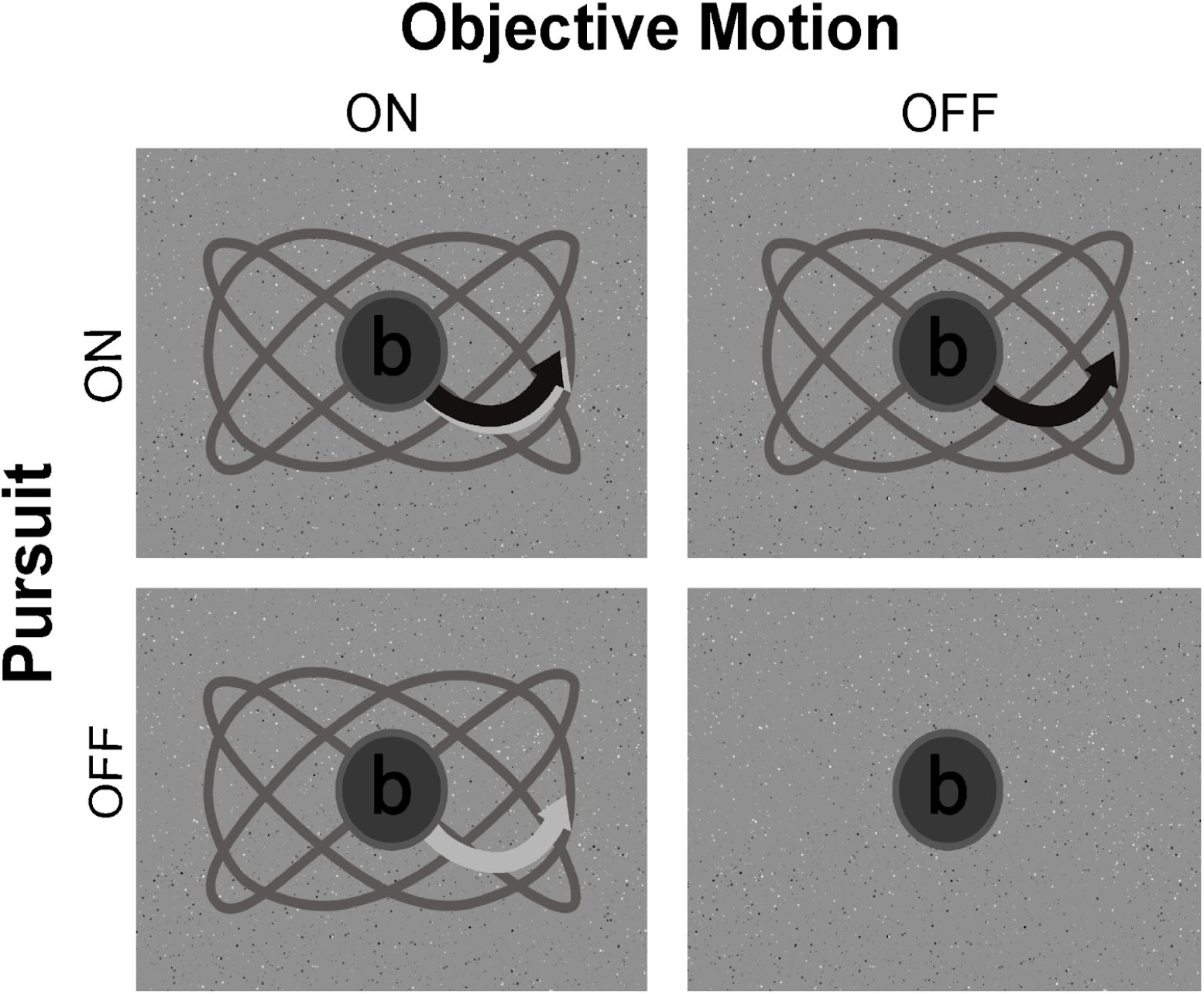
Schematic illustration of stimuli and 2×2 factorial design. The two factors were pursuit (on/off) and objective motion (on/off). In pursuit ‘on’ conditions, the fixation-task moved on the screen on a sinusoidal path (black arrow; grey path), with a maximal eccentricity of 2.5 degrees. In the objective motion ‘on’ conditions, the dot-field moved along the same trajectory (light grey arrow; grey path). Trajectories varied randomly across trials. A 1-back character matching task was presented on the fixation disk at all times. Note that the contrast for objective motion (left column versus right column) and the contrast for retinal motion (ascending diagonal versus descending diagonal) were both fully controlled, as peripheral effects induced by pursuit canceled each other out within each contrast. Fixation and trajectory are enlarged for visualization; actual trajectory reached 2.5 degrees eccentricity, actual screen size was 30×20 degrees.

Two key contrasts available from this 2×2 design, namely the contrast “objective motion” (i.e. both conditions with objective motion “on” versus both conditions with objective motion “off”), and “retinal motion” (i.e. the two conditions where pursuit and objective motion were in different states (on/off and off/on) and the two conditions where both factors were in the same state (on/on and off/off)) were fully controlled for the amount of retinal motion and pursuit signals. As each of these contrasts had one pursuit condition counting positive and one counting negative, any peripheral motion artifacts resulting from pursuit cancelled each other out within each contrast. Hence, foveal and parafoveal representations (out to 12.5 × 7.5 visual degrees eccentricity) would be stimulated in a fully controlled way, and peripheral effects would be balanced and cancel out. Only the contrast “pursuit” (both pursuit “on” conditions versus both pursuit “off” conditions) was difficult to interpret as it combined motor effects of pursuit with visual effects (peripheral motion), which is why we focus here on the first two, controlled, contrasts, as we did in our original study (Fischer et al., 2012).

### Procedure

Stimulus presentation followed a block design schedule where each of the four stimulus conditions was presented eight times with a duration of 12 s each. Hence, each of the four functional runs consisted of 32 full stimulus blocks. Different conditions were presented in pseudorandom sequences in which each condition was preceded equally often by all conditions. The stimulus was rear-projected using a linearized projector with a resolution of 1024 × 768 pixels at 60 Hz onto a screen located in the scanner bore. Subjects viewed the stimulus at 82 cm distance through a mirror mounted on the receive coil array, leading to a display size of 30 × 20 visual degrees. The stimulus was written in MATLAB 2010a (http://www.mathworks.de/) using the Psychophysics Toolbox 3 extensions (http://psychtoolbox.org/) and was presented using a Windows computer.

### Fixation Task

Throughout the experiment subjects performed a character repetition-detection task on a fixation disc, ensuring fixation as well as balanced attention across conditions. A total of 26 characters were presented in random succession (1.6 degrees height, white) on a gray fixation annulus (2 degrees width, 72 cd/m2), with random presentation times of 1-2.16 s. Subjects indicated character repetitions by button press.

### Data Acquisition and Image Reconstruction

All measurements were conducted on a 9.4 Tesla whole-body MRI scanner (Siemens, Erlangen, Germany) using a custom-built head coil with a 16-element dual row transmit array and a 31-element receive array (Shajan et al., 2014).

For the acquisition of blood oxygen level dependent (BOLD) weighted images, we used PSF (Point Spread Function) corrected (In and Speck, 2012) 2D gradient-echo EPI with 0.8 × 0.8 × 0.8 mm isotropic resolution and the following parameters (see section below for PSF correction details): TR / TE / flip angle = 2000 ms / 21 ms / 70 deg, field of view = 160 × 160 mm, matrix = 200 × 200, bandwidth = 1388 Hz / pixel, GRAPPA (Griswold et al., 2002) acceleration factor of 4, and partial Fourier of 6 / 8. The slices covered the dorsal part of the visual cortex and part of parietal cortex using 20 oblique-coronal slices positioned parallel to the calcarine sulcus. The number of slices was limited to 20 due to limits in the specific absorption rate (SAR). The wall-time for each run was 6 min and 36 seconds (396 s), including the initial 8 s block of dummy scans in the beginning of each scan to allow T1 steady state to be achieved, and 1 TR of reference scan. Functional images were reconstructed with the standard online Siemens EPI and GRAPPA reconstruction. Four functional runs each consisting of 203 volumes, including the four dummy scans and one reference scan, were obtained for each subject. For each run, automatic reconstruction of data acquisitions was done twice, once with PSF correction and once without PSF correction. PSF corrected images were used for the subsequent analysis.

Whole-brain T1-weighted anatomical images were acquired for each subject using a MP2RAGE sequence (Marques et al., 2010) (TR = 6 ms, TE = 2.3 ms, voxel size 0.8 × 0.8 × 0.8 mm, matrix = 256 × 256 × 192, PAT mode: GRAPPA 3, and 6/8 partial Fourier with POCS reconstruction), yielding two inversion contrasts (flip angle 1 = 4°, flip angle 2 = 6°, TI1= 900 ms, TI2 = 3500 ms). MP2RAGE data was reconstructed offline using custom software developed in MATLAB.

### Preprocessing and Statistical Analysis of Functional Volumes

#### Point-Spread Function Correction (PSF) of the EPI Images

Point-spread function (PSF) mapping is one of the promising methods for correcting geometrical and intensity-related distortions in Echo-planar imaging (EPI). Using acquisitions with additional phase-encoding gradients, PSF maps encode spatial information relevant to overall intensity distribution and geometrical distortion from a single voxel. These maps are then convolved with the distorted image in order to obtain PSF-corrected EPIs (Zeng and Constable, 2002). We used PSF mapping to correct for (1): possible distortions in the EPI images that are related to field inhomogeneity, (2): eddy current effects, and (3): blurring due to image distortion. Pixel shifts in image space were corrected based on a PSF map (Zeng and Constable, 2002; Zaitsev et al., 2004; In and Speck, 2012). The point spread function was measured in a separate scan prior to each BOLD sequence using the same parameters as the EPI sequence. PSF shift maps in the non-distorted spin-wrap encoding direction (obtained online) were used to compute the average and the standard deviation of the shift values in the regions-of-interest (ROIs). Compared to distorted images, PSF-corrected activation maps were registered easier and with more precision to the anatomical images.

#### Preprocessing and GLM analysis

PSF corrected functional data were preprocessed and analyzed using the FreeSurfer functional analysis stream, FSFAST (https://surfer.nmr.mgh.harvard.edu/fswiki/FsFast). Preprocessing included motion correction of the functional volumes to the first volume of the first run, slice-timing correction, and co-registering functional images to the MP2RAGE. No spatial smoothing (volume- or surface-based) was applied to the data at any stage of the analysis.

A general linear model (GLM) including regressors for each condition as well as head-motion parameters was then fitted to the time course of each voxel. A second order polynomial function was used as a nuisance regressor to model low frequency drifts. The GLM analysis of the 2×2 factorial design allowed us to separate cortical responses related to the main factors of (a) eye-movements (active pursuit), (b) objective (2D planar) motion, and their interaction: (c) retinal motion. The main factor (b: 2D planar motion) and the interaction (c: retinal motion) were balanced for all effects of pursuit, hence cancelling effects related to motion of the screen edges induced by eye movements or to less accurate fixation during pursuit.

#### Definition of Regions of Interest

FreeSurfer surface-based analysis (Dale et al., 1999; Fischl et al., 1999; Polimeni et al., 2018) was used to define the ROIs on the surface. For ROI definition only, fMRI data was smoothed (3 mm) on the surface – note that for ROI definition, neither cortical depth information nor ultra-high resolution were relevant. No smoothing, neither in the volume nor on the surface, was performed in any stage of the subsequent laminar analysis. We used a previously established motion localizer to localize V3A and V6+ (Fischer et al., 2012) as follows. Using the contrast ‘objective versus retinal motion’, V3A and V6+ were defined as voxels that had higher responses to objective (‘real’) compared to retinal motion and that were located in the anatomical regions consistent with prior studies. In these prior studies, this contrast has led to selective activation of voxels overlapping with retinotopically defined V3A (Fischer et al., 2012). V3A was located below the parietal-occipital sulcus (POS) and extended into the transverse occipital sulcus (TOS) coinciding with the anatomical landmarks previously reported for V3A (Tootell et al., 1997; Silver et al., 2005; Pitzalis et al., 2006). The same functional contrast was previously also shown to activate voxels overlapping retinotopically defined V6 (Fischer et al., 2012), which is located in the dorsal part of the POS (Pitzalis et al., 2006, 2015). However, for V6 we cannot exclude the possibility that the neighboring retinotopic area V6A responded also to our localizer contrast. Recent studies reported that V6A also responded to visual flow fields (Pitzalis et al., 2013, 2015). We hence refer to this functionally defined region as V6+ to indicate that it includes V6 and possibly V6A.

In every subject the first functional run was used to localize V3A and V6+, and the remaining three runs were used for analysis.

In addition to investigating responses in V3A and V6+, which were the main focus of this study, we also identified area V7 (also referred to as IPS-0) (Tootell et al., 1998). V7 shares a common feature with V3A and V6+, in that it, too, responds to coherent motion (Cardin and Smith, 2011; Helfrich et al., 2013) and eye movements (Schluppeck et al., 2005). However, compared to V3A and V6+, V7 has no or only small preference to objective motion compared to retinal motion (Fischer et al., 2012; Nau et al., 2018). As it is also directly neighboring V3A and V6+, and as it is located within the narrow field of view of our focused imaging sequence, we selected V7 as a control ROI to examine to which extent the pattern of results we find in V3A and V6+ can be attributed to their overall preference to objective motion. To localize V7 we used maximum probability maps (most probable region for any given point on each individual’s cortical surface) of the functional probabilistic atlas provided by the Kastner group (Wang et al., 2015), thresholded at 50 percent.

#### Depth Dependent Analysis

FreeSurfer (Dale et al., 1999) was used to generate surface reconstructions of the interface between white matter and gray matter (white surface) and between gray matter and CSF (pial surface) from 0.8 mm MP2RAGE data, and to create cortical thickness maps from these boundaries (Fischl and Dale, 2000), using a modified reconstruction stream adapted for MP2RAGE (Fujimoto et al., 2014). Following Polimeni et al. (2010), nine additional surfaces within gray matter at fixed relative distances from white matter and pial surface were then derived from cortical thickness maps (Polimeni et al., 2010). Thus, in total eleven surfaces corresponding to white matter surface, pial surface, and nine equally spaced intermediate surfaces were created for each subject and each hemisphere (Figure 2).

**Figure 2:**
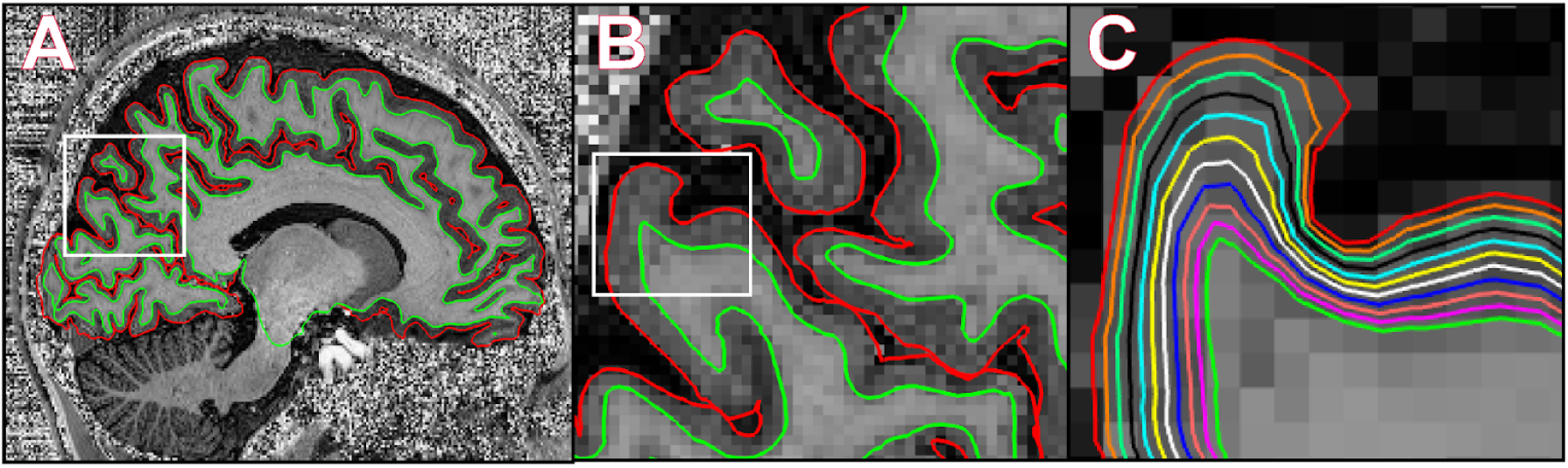
Laminar surface segmentation. (A) A T1w MP2RAGE image was used to segment the brain and to generate intermediate surfaces throughout the whole cortex. (B) The approximate location of V3A and V6+ is magnified to illustrate the sections shown in (C). (C) Illustration of laminar depth-segmentation using nine surfaces between white-gray matter boundary and pial surface.

Functional volumes were aligned to the surface reconstructions generated from the MP2RAGE anatomical data using a boundary-based registration method (Greve and Fischl, 2009) that first identified the WM-GM boundary in the EPI data and then registered this interface to the corresponding surface reconstruction in the anatomical image using a rigid transformation. To map functional signal to the surface reconstructions we resampled GLM statistics of each voxel in the functional volume to the eleven surfaces reconstructed from the anatomical MP2RAGE image by nearest neighbor interpolation. When the size of the fMRI voxel is greater than the spacing between mesh vertices, which is the case in this study, nearest neighbor interpolation has been shown to work well in mapping the functional signal onto the surface reconstructions (Polimeni et al., 2018).

#### Statistical Analysis

Laminar signal was analyzed in three ways for each ROI. First, a two-way ANOVA compared signal intensities as a function of cortical depth (factor depth) and of motion type (factor motion type: retinal motion, objective motion) using the GLM-derived contrast values.

Second, given the fact that contrast estimates for different signal types (retinal, objective) showed an increase from lower towards upper layers, we examined whether the slopes of this increase differed between signal types. The slopes were calculated by fitting a first degree polynomial function to eleven data points corresponding to the eleven laminar signals.

Third, we performed the above-described two-way ANOVA after detrending signal intensities across depth: given the known signal bias towards larger vessels in GE-EPI, signal increases towards superficial layers, and differences in signal slopes could be due to differential blood vessel distribution across the cortical depth and differences in proximity to the large surface vessels, rather than due to neural effects (or in addition to the latter). In an attempt to overcome this surface bias, we repeated the two-way ANOVA following linear detrending of laminar responses of retinal and objective motion. Using linear regression, we removed constant and linear trends from the laminar profiles of each motion type and examined only the residual effects, now corrected for the overall linear signal increase across layers and for differences in mean signal amplitude. The analysis of the residuals would therefore reveal the uniqueness of condition-specific laminar profiles beyond surface-bias and mean differences.

## RESULTS

We used ultra-high field (9.4T) fMRI to examine laminar activation profiles of high level visual areas V3A, V6+, and V7 in response to retinal and objective motion during pursuit eye movement in eleven human participants.

In the separate localizer runs, we were able to reproduce the previously reported response to objective and to retinal motion in V3A and V6 (Fischer et al., 2012). Every subject revealed clusters with objective motion preference in anatomical positions corresponding to typical locations of V3A and V6+, as shown in Figure 3 for three representative subjects.

**Figure 3:**
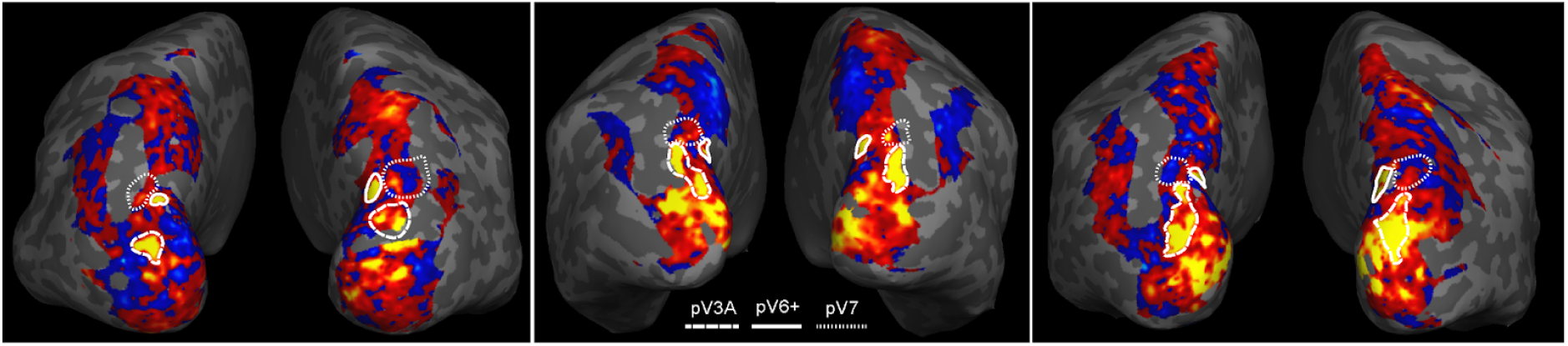
Cortical responses to objective and retinal motion and boundaries of putative V3A, V6+ and V7 on representative subjects. For better visualization and more accurate estimation of ROI boundaries, smoothed (3mm) activations were used. Cortical surface that was covered by our high-resolution functional slab is colored. For putative V3A and V6+, the labelled outlines show objective motion responsive patches overlapping with probabilistic maps (Wang et al., 2015) (for V3A) and overlapping with the typical position in the superior posterior occipital sulcus (for V6+); for putative V7, the outlines represent the probabilistically mapped V7 (see methods, “Definition of Regions of Interest” section for details).

### Laminar Responses to Retinal and Objective Motion in V3A and V6+

In both V3A and V6+, the average net BOLD signal was higher during objective motion compared to retinal motion (V3A: t(21)=-8.47, p=3.24*10^−8^, V6+: t(21)=-6.57, p=1.64*10^−6^). Like all results shown below, this result is based on the last three imaging runs and hence confirms the results from the first run that was used to select the ROIs based on this contrast.

The laminar profiles for retinal and objective motion differed in both ROIs, V3A and V6+, as is evident in Figure 4 (A, B). The two-way ANOVA showed that there was a significant interaction between cortical depth and motion type in both regions (V3A: F(10,210)=22.59, p<0.001; V6+: F(10,210)=20.18, p<0.001). For V3A, post-hoc paired sample t-test across depth levels revealed a significantly stronger response to objective compared to retinal motion in the superficial layers, but not in the lower layers (upper layers: depths 9 and higher: t(21)< -3.11, p<0.0053; lower layers: depth 1: t(21)=-0.59, p=0.56, depth 2: t(21)=-0.95, p=0.35, depth 3: t(21)=-1.25, p=0.23) (Figure 4A). In V6+ the response to objective motion was stronger than that to retinal motion across all depths, yet with larger differences towards the superficial layers (Figure 4B). These results were also reflected in the slopes analysis, that showed significantly higher slopes in the objective motion compared to the retinal motion in both regions (V3A: t(21)=-6.32, p=2.0*10^−6^, V6+: t(21)=-5.66, p=7.0*10^−6^).

**Figure 4:**
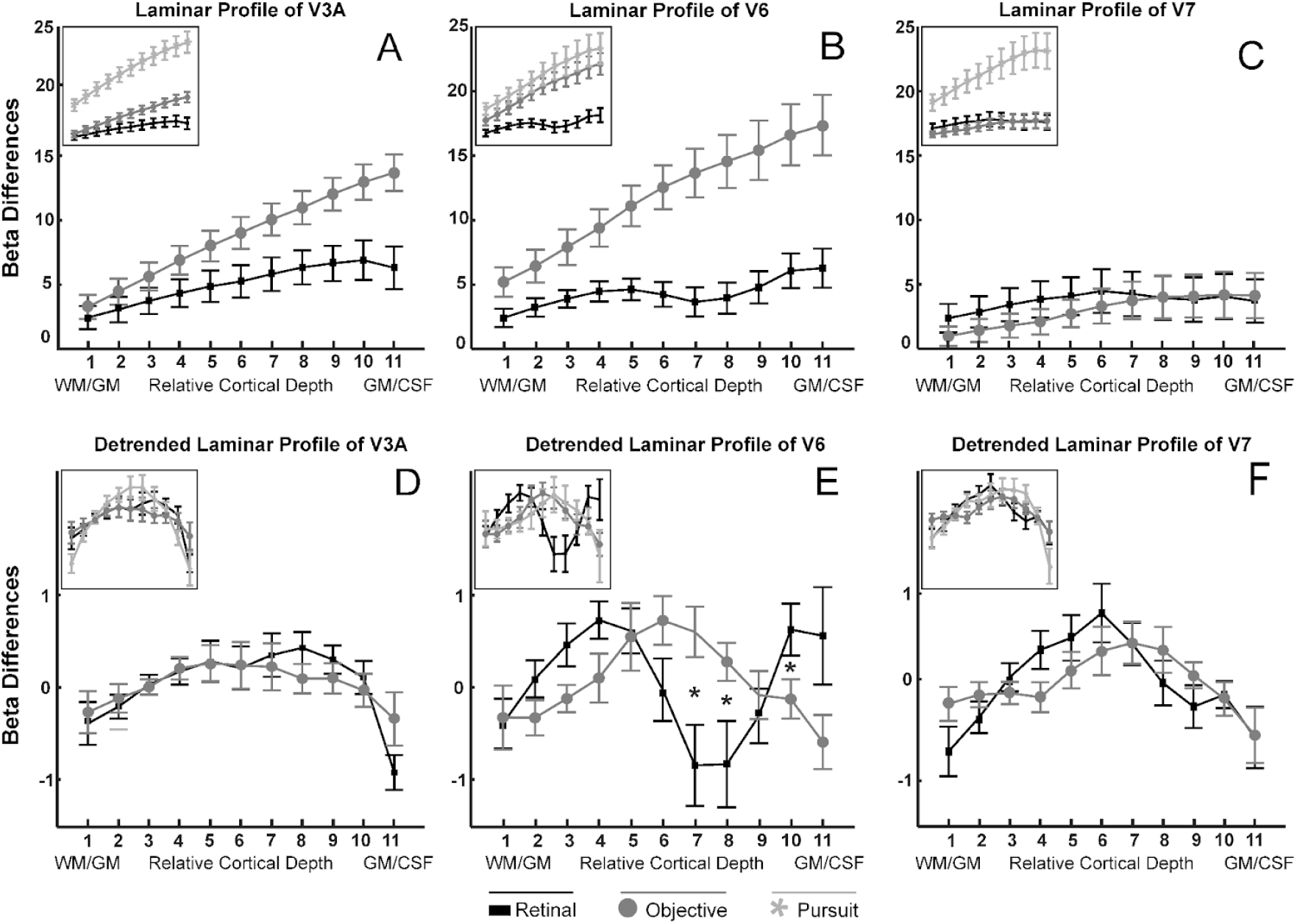
Laminar profile of putative V3A, V6+, and V7 in response to retinal and objective motion. (A, B, C): raw laminar signals in response to the motion types, (D, E, F): laminar signals from which the linear trend and the overall mean signal were removed. The insets additionally show responses to the pursuit contrast, which was not further considered as it confounds peripheral visual motion with other aspects of pursuit. Two-way ANOVAs (factors: depth, motion type) showed significant interactions between depth and motion type in the raw depth profiles (A-C) in V3A, V6+, and V7 (V3A: F(10,210)=22.59, p<0.001; V6+: F(10,210)=20.18, p<0.001; V7: F(10,210)=2.62, p=0.0051). For the detrended depth profiles (D-F), the ANOVAs revealed a significant interaction in V6+ (V3A: F(10,210)=1.59, p=0.052; V6+: F(10,210)=2.07, p=0.0045). Error bars represent SEM across subjects. Post-hoc t-tests were conducted at each depth level between values of retinal and objective motion. *: p<0.05 uncorrected.

We included V7 as a control ROI in order to test whether the upper-layer bias for objective motion could also be observed in a region that had no overall functional preference for objective motion (Fischer et al., 2012; Nau et al., 2018). Hence, if V7 showed an interaction between motion type and depth similar to V3A and V6+, this would speak for a genuine neural bias for objective motion in the upper layers.

Consistent with previous results, V7 showed no preference for objective compared to retinal motion (Figure 4C). Even though weak, the laminar results for V7 support the neural bias account. The ANOVA showed a significant interaction between layer and motion type (Figure 4C) (F(10,210)=2.62, p=0.0051), which was driven by a marginally steeper laminar response slope in V7 for objective compared to retinal motion (slope analysis: t(21)=-1.83, p=0.082). Note that this result cannot be accounted for by overall higher responses to objective motion, as, to the contrary, retinal signal trended higher than objective motion in lower layers (p<0.2 at each layer). The upper-layer response bias during objective compared to retinal motion processing in V3A and V7 hence lend support to the notion of enhanced neural activity in upper layers during objective motion processing.

### Detrended Laminar Response profiles in V3A and V6+

Since all responses contained activity biases towards upper layers we wanted to perform an additional analysis that examined the underlying laminar response profiles after removal of each individual layer bias from each ROI and each condition.

Figure 4 (D, E, F) shows the laminar profiles of each ROI and each motion type after removal of the linear trend and the overall mean signal. A prominent dip in V6+’s mid-layer retinal motion response is apparent. This is reflected by a significant interaction between depth and motion type in V6+ in the two-way ANOVAs (V6+: F(10,210)=2.07, p=0.0045; V3A: F(10,210)=1.59, p=0.0518). Per-depth post-hoc paired-sample t-tests in each ROI revealed significant differences between laminar profiles of retinal and objective motion only in V6+ at depths 7, 8, and 10 (depth 7: t(21)=-2.62, p=0.016; depth 8: t(21)=-2.39, p=0.026; depth10: t(21)=2.17, p=0.042). In order to test additionally whether the depth profiles of the motion types (retinal or objective motion) differed between ROIs, we carried out a second two-way ANOVA with the factors depth and ROI (two ROIs in each ANOVA), separately for retinal and for objective motion. This ANOVA revealed that retinal motion in V6+ differed from that in V3A and V7 (V6+ and V3A: F(10,210)=4.02, p=0.5*10^−6^; V6+ and V7: F(10,210)=2.9, p=0.002) but that objective motion did not differ between ROIs (V6+ and V3A: F(10,210)=0.86, p=0.57; V6+ and V7: F(10,210)=0.37, p=0.96).

Figure 5 shows the superimposed depth profiles of retinal motion for V6+ and V3A, and for V6+ and V7, along with post-hoc per-depth t-tests that again highlight the mid-layer dip in V6+ (V6+ and V3A: depth 4: t(21)=-2.61, p=0.016; depth 7: t(21)=2.52, p=0.020; depth 8: t(21)=2.68, p=0.014; depth 9 t(21)=2.19, p=0.040; depth 11 t(21)=-2.49, p=0.021; V6+ and V7: depth 7: t(21)=-2.57, p=0.018).

**Figure 5:**
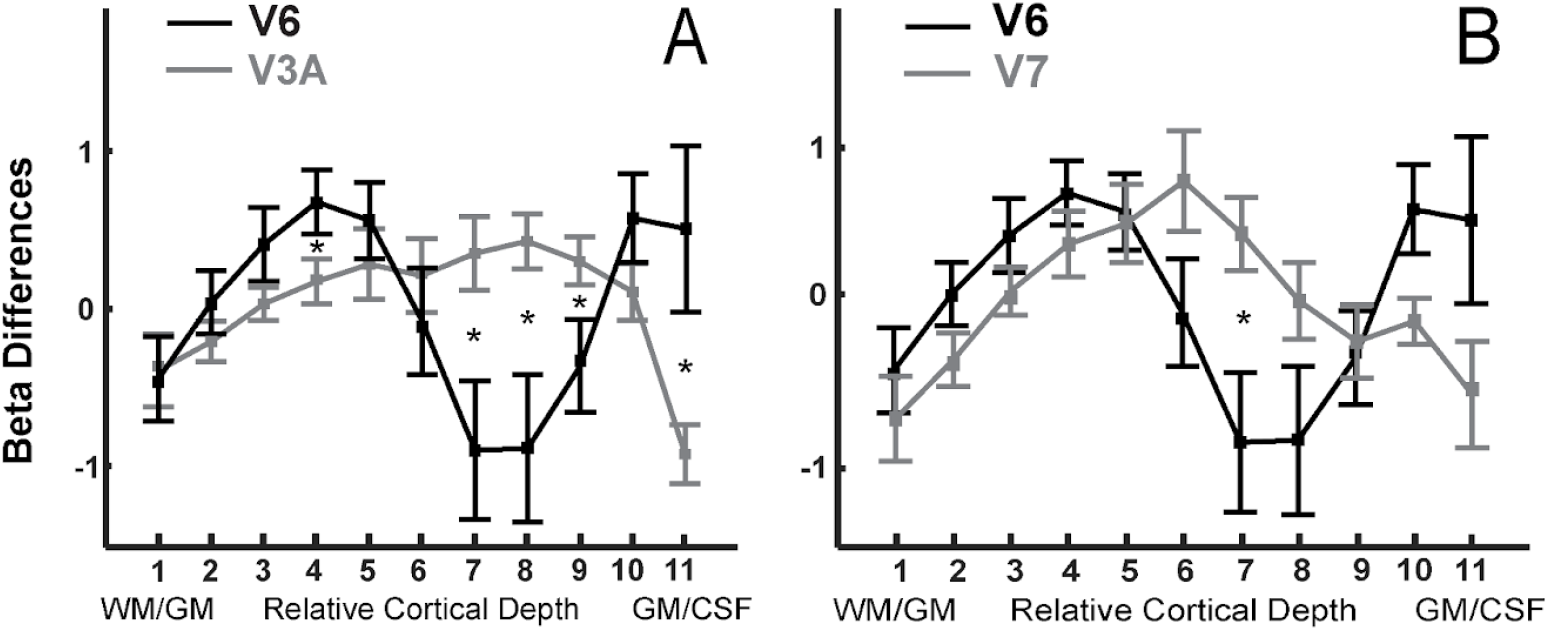
Superimposed depth profiles of retinal motion for V6+ and V3A, and for V6+ and V7. The mid-layer dip in V6+, along with the significant difference between depth profiles of retinal motion in V6+ compared to V3A and V7, is highlighted by post-hoc per-depth t-tests (V6+ and V3A: depth 4: t(21)=-2.61, p=0.016; depth 7: t(21)=2.52, p=0.020; depth 8: t(21)=2.68, p=0.014; depth 9 t(21)=2.19, p=0.040; depth 11 t(21)=-2.49, p=0.021; V6+ and V7: depth 7: t(21)=-2.57, p=0.018). *: p<0.05 uncorrected.

Together, the ANOVA results show that the depth profile for retinal motion in V6+ was unique in that it differed from that of objective motion, as well as from retinal motion profiles of neighboring regions V3A and V7. In contrast, the depth profiles for objective motion did not differ across regions.

## DISCUSSION

In this study, we used ultra-high field (9.4T) fMRI to measure laminar BOLD signal in high-level visual areas V3A, V6+ and V7 in response to retinal motion, objective motion, and pursuit eye-movement. As estimation of objective motion involves a multi-modal comparison between (visual) retinal motion signals and (non-visual) efference copies, we hypothesized differential processing across cortical depth associated to objective motion compared to retinal motion. In order to test this, we used a factorial design including objective motion and smooth pursuit eye movements that has previously been shown to allow functional segregation of retinal and objective motion responses (Fischer et al., 2012).

Our results firstly demonstrate that laminar signals at high-level motion regions such as V3A, V6+, and V7 can be captured using ultra-high field fMRI, and we confirmed a preference of objective over retinal motion in V3A and V6. Laminar analysis showed an interaction between depth and experimental condition in all regions, driven by a stronger signal increase towards superficial layers for objective compared to retinal motion. In V3A, this was the case even though signal strength in lower layers was matched. Finally, we uncovered an area- and motion-type specific differential depth profile in V6+ after removing the upper-layer bias, revealing a mid-layer dip for retinal motion not present in other areas or for objective motion.

### Laminar Profiles

There are two possible accounts for the upper-layer bias results obtained using raw signal (Figure 4 A-C). First, the measured response profiles are compatible with segregated neural responses across depth, namely a higher fraction of voxels with a preference towards objective motion in superficial layers. This population of voxels could be driven by feedback signals related to pursuit and/or by local processing integrating pursuit signal with retinal signal entering at middle layers. In the context of this study we speculate that these feedback projections transmit efference copies of the eye movement from other cortical areas, potentially originating from pursuit regions of frontal eye fields (Stanton et al., 2005) or MST (Boussaoud et al., 1990), to superficial layers in V3A and V6+.

Second, as the laminar responses did not show a double-dissociation between experimental conditions across depth but instead an increase of objective motion response towards upper layers only, an alternative account driven by pial vessels and diving venules inducing a spatial bias of increased BOLD signal toward superficial layers (Ahveninen et al., 2016; Polimeni et al., 2018) cannot be excluded. This is especially so as V3A and V6+ were localized according to their higher preference to objective compared to retinal motion, with stronger signal potentially associated to stronger upper-layer bias.

Two observations speaking against this account are the lack of signal differences in lower layers in V3A, and the higher slope for objective motion in control area V7. Area V7 does not have an overall objective motion bias, and, similar to V3A and MST, is involved in eye-movement related processing (Schluppeck et al., 2005; Konen and Kastner, 2008). The laminar response profile in V7, just as those of V3A and V6+, are hence consistent with the arrival of pursuit related feedback signals in the superficial layers.

### Detrended Laminar Profiles

All detrended laminar profiles showed a higher response in middle layers surrounded by lower responses in deep and superficial layers. The only exception was the laminar profile in V6+ to retinal motion that showed a prominent dip in the middle layers. The present results open up several interesting mechanistic interpretations that will need to be investigated in future studies.

First, we know that objective motion responses are the product of a subtractive comparison between pursuit signals and retinal motion, and that in 3T fMRI, V6 was suppressed by retinal motion alone (Fischer et al., 2012). Reduction of activity in middle layers of V6+ can be a manifestation of this subtractive comparison mechanism, which cancels retinal motion by silencing the retinal motion signals entering V6+. This mechanism could account for the previously observed suppression of V6 by retinal motion (Fischer et al., 2012).

Alternatively, the mid-layer dip in V6+’s retinal motion profile can be seen as relative enhancement in deep and superficial layers, indicative of increased descending inputs coming from higher-level regions, such as MST and entering extragranular layers of V6+. This is consistent with the known laminar profiles of descending neuroanatomical connections (Felleman and Van Essen, 1991), and would be supported by the extensive connectivity of V6 with other motion responsive regions (Galletti et al., 2001; Galletti and Fattori, 2003; Fischer et al., 2012; Galletti and Fattori, 2018). This view is also compatible with the large receptive fields in V6 and its preference for complex ego-motion compatible motion (Pitzalis et al., 2006, 2010, 2012, 2015; Cardin and Smith, 2010, 2011), and its potential role in complex vision-for-action tasks (Galletti et al., 2003, 2018; Pitzalis et al., 2013, 2015).

While future studies are needed to differentiate between the different interpretations, to our knowledge, this has been the first study demonstrating condition-specific laminar profile differences in a high-level visual motion region, providing a new entry to a better mechanistic understanding of motion processing mechanisms in V3A and V6+.

### Laminar Organization of Neuronal Activations

In the sensory system, bottom-up feedforward signals convey sensory inputs from the external world into the brain, whereas top-down feedback signals are thought to project higher-level information such as expectations or non-visual information to visual regions. Feedforward and feedback signals are largely segregated and terminate in distinct layers of the cortex (Wong-Riley, 1978; Felleman and Van Essen, 1991; Petro and Muckli, 2017). Contextual BOLD effects in superficial layers can also arise from layers II and III, known to be targeted by horizontal connections (Angelucci and Bullier, 2003; Self et al., 2013).

One possible mechanism to explain how interaction with bottom-up sensory inputs can change the laminar profile of feedback signals is through inhibitory connections from the deep layers to the granular layer IV (Thomson and Bannister, 2003; Katzel et al., 2011; Kim et al., 2014), which leads to a reduction throughout the entire cortical column as a result of the excitatory pathway from layer IV to layers II–III and from layers II–III to layers V–VI (Douglas and Martin, 2004). It should, however, be noted that local excitatory-inhibitory circuitries may have distinct characteristics across different cortical regions and also across different species. Therefore, the mechanism of integration described here may not be fully translatable to V3A and V6+ areas in human.

It has previously been shown that remapping of the eye movement motor commands in V3A precedes occurrence of the eye movement (Nakamura and Colby, 2002) suggesting that this area has access to the information about the eye movement before efference copies of the eye movement reach them. This could suggest that the observed laminar profile of V3A may be driven, at least in part, by the arrival of efference copies.

### Limitations of Laminar Imaging

Venous blood draining from deep layers toward superficial layers pools deoxygenated blood towards the surface (Duvernoy et al., 1981; Chen et al., 2013). Therefore, the closer to the cortical surface the more laminar activation is expected to contain a mixture of signals from lower layers. These laminar interdependencies make it difficult to precisely ascertain the origin of laminar activation, especially in the superficial layers.

Methodological constraints of cortical depth sampling are another important consideration in laminar imaging. For instance, relative thickness of cortical laminae changes across the cortex and correlates with the local folding pattern (Van Essen and Maunsell, 1980; Fatterpekar et al., 2003). Therefore, depths sampled on equally distanced cortical positions based on normalized distances between the white matter and pial surface boundary (Dale et al., 1999; Fischl et al., 1999, 2000, 2001, 2002; Segonne et al., 2004; Polimeni et al., 2010), do not precisely correspond to the histological laminar anatomy per se. Although such surface reconstructions are not expected to have a perfect alignment with cortical lamina, many high-field laminar studies (Koopmans et al., 2010, 2011; Polimeni et al., 2010; Zimmermann et al., 2011; Olman et al., 2012; De Martino et al., 2015) have demonstrated the robustness of this approach in revealing functional response profiles across the cortical depth with high resolution functional data.

Finally, neurovascular coupling can differ for negative and positive BOLD responses as well as across layers (Goense et al., 2012). However, studies in macaques have shown that neuronal excitation (Goense and Logothetis, 2008) and inhibition (Shmuel et al., 2006) typically closely correspond to the BOLD signal, even though exceptions do exist (Bartels et al., 2008; Logothetis, 2008).

Our study demonstrates the ability of high field fMRI to capture neuronal activation at the level of fine scaled cortical structures i.e. cortical layers. Investigation of the information content of laminar signals (Kamitani and Tong, 2005; Williams et al., 2008; Muckli et al., 2015; Klein et al., 2018) rather than their amplitude can alternatively be pursued in future studies to investigate other aspects of laminar signals.

## Acknowledgements

This study was funded by the Center for Integrative Neuroscience Tübingen (the German Excellence Initiative of the DFG, grant number EXC307), and by the Max-Planck Society, Germany. We thank Dr. Andreas Schindler for his help in implementing visual stimulation.

